# Unraveling ChR2-driven stochastic Ca^2+^ dynamics in astrocytes – A call for new interventional paradigms

**DOI:** 10.1101/549469

**Authors:** Arash Moshkforoush, Lakshmini Balachandar, Carolina Moncion, Josue Santana, Jorge Riera Diaz

## Abstract

Control of astrocytes via modulation of Ca^2+^ oscillations using techniques like optogenetics can prove to be crucial in therapeutic intervention of a variety of neurological disorders. However, a systematic study quantifying the effect of optogenetic stimulation in astrocytes is yet to be performed. Here, we propose a novel stochastic Ca^2+^dynamics model that incorporates the light sensitive component – channelrhodopsin 2 (ChR2). Utilizing this model, we studied the effect of various pulsed light stimulation paradigms on astrocytes for select variants of ChR2 (wild type, ChETA, and ChRET/TC) in both an individual and a network of cells. Our results exhibited a consistent pattern of Ca^2+^ activity among individual cells in response to optogenetic stimulation, i.e., showing steady state regimes with increased Ca^2+^ basal level and Ca^2+^ spiking probability. Furthermore, we performed a global sensitivity analysis to assess the effect of stochasticity and variation of model parameters on astrocytic Ca^2+^ dynamics in the presence and absence of light stimulation, respectively. Results indicated that directing variants towards the first open state of the photo-cycle of ChR2 (o_1_) enhances spiking activity in astrocytes during optical stimulation. Evaluation of the effect of astrocytic ChR2 expression (heterogeneity) on Ca^2+^ signaling revealed that the optimal stimulation paradigm of a network does not necessarily coincide with that of an individual cell. Simulation for ChETA-incorporated astrocytes suggest that maximal activity of a single cell reduced the spiking probability of the network of astrocytes at higher degrees of ChR2 expression efficiency due to an elevation of basal Ca^2+^ beyond physiological levels. Collectively, the framework presented in this study provides valuable information for the selection of light stimulation paradigms that elicit optimal astrocytic activity using existing ChR2 constructs, as well as aids in the engineering of future optogenetic constructs.

**Author summary:** Optogenetics – an avant-garde technique involves targeted delivery of light sensitive ion channels to cells. Channelrhodopsin 2 (ChR2), an algal derived light sensitive ion channel has extensively been used in neuroscience to manipulate various cell types in a guided and controlled manner. Despite being predominantly used in neurons, recent advancements have led to the expansion of the application of optogenetics in non-neuronal cell types, like astrocytes. These cells play a key role in various aspects of the central nervous system and alteration of their signaling is associated with various disorders, including epilepsy, stroke and Alzheimer’s disease. Hence, invaluable information for therapeutic intervention can be obtained from using optogenetics to regulate astrocytic activity in a strategic manner. Here, we propose a novel computational model to assess astrocytic response to optogenetic stimulation which implicitly accounts for the stochastic character of Ca^2+^ signaling in this cell type. We identified light stimulation paradigms suitable for eliciting astrocytic Ca^2+^ response within physiological levels in widely-used ChR2 variants and identified highly sensitive parameters in ChR2 kinetics conducive for higher probability in Ca^2+^ spiking. Overall, the results of this model can be used to boost astrocyte light-induced behavior prediction and the development of improved future optogenetic constructs.

## Introduction

Astrocytes - key players in the brain, are involved in neurovascular coupling [1–3], serve as communication elements and regulate neuronal activity via gliotransmission [4–6]. They are pivotal in housekeeping roles such as providing metabolic support to neurons [7, 8], rendering cytoarchitectonic support to the brain environment, and maintaining carbon homeostasis which leads to the regulation of ‘excitatory – inhibitory’ neurotransmitter balance [9]. Astrocytes are extensively involved in the reduction of toxicity in the neuronal environment through scavenging reactive oxygen species, thereby minimizing tissue damage [10]. In the case of neurotoxic insults, they assist microglia in the *de-novo* synthesis of various cytokines and trophic factors resulting in the modulation of neuroinflammation [11–13]. Dysregulation of astrocytic function results in a multitude of brain disorders including epilepsy, stroke and Alzheimer’s disease [14–20]. Hence, control of astrocytes is a powerful tool for intervening and preventing brain dysfunction. Since calcium signaling is one of the major regulatory mechanisms in astrocytes, its control can serve as a target for therapeutic intervention [21–25].

Several research groups have demonstrated the ability to elevating Ca^2+^ activity in astrocytes via electrical [26–29], mechanical [30–32] and pharmacological [33, 34] approaches. Upon electrical stimulation, astrocytes exhibit high frequency oscillations, mainly through L-type Ca^2+^ channels. However, this methodology lacks cell specificity due to potential concurrent activation of neurons and suffers low spatial resolution. Additionally, the feasibility of this method has not yet been tested *in vivo*. Mechanical stimulation, performed to mimic responses to brain injury and spreading depression [29, 35], lacks clinical feasibility. The use of pharmacological techniques for targeting these cells in the brain has been limited to basic research due to high invasiveness and low temporal resolution [36, 37]. Contrarily, optogenetics is an avant-garde minimally invasive approach, which in combination with advancements in the field of nonlinear optics [38–40], has provided a platform for genetically targeting specific cell types with high temporal and spatial precision [37, 41–43].

In spite of the recent inception of the field of optogenetics, a wide variety of optogenetic tools have been constructed, among which channelrhodopsin 2 (ChR2) has been one of the most commonly used. There exists an extensive body of literature on the biophysical characterization of ChR2 variants and their response to various light stimulation paradigms, predominantly in excitable cells [44–47]. For example, many research groups have engineered ChR2 variants for enhanced conductance, increasing recovery kinetics and capability of stimulation at lower light levels in neurons [44, 48]. ChR2 variants have also been modified to form chimeric variants for regulating responses and facilitating multiwavelength optogenetics in neurons [49]. There have been few studies on optogenetically targeting astrocytes for specific applications [50–54], including their role in memory enhancement [55] and cortical state switching [56]. However, a holistic approach to quantify the effect of light stimulation on astrocytes has not yet been formulated, a vital step for strategic manipulation of these cells. In analyzing this effect, accounting for the stochastic nature of spontaneous calcium oscillations (SCOs) in astrocytes is imperative. The source of this stochasticity is primarily ascribed to the randomness in fluxes through IP_3_R clusters and the plasma membrane (PM) [57, 58].

This paper seeks to provide a comprehensive platform via mathematical modeling to optimize light stimulation paradigms for existing optogenetic variants, yielding high astrocytic spiking rates without eliciting non-physiological behavior, and to aid the development of novel application-based constructs targeting astrocytes. To this end, we outline a novel stochastic model of astrocyte calcium dynamics with an incorporated optogenetic component - ChR2. Firstly, we quantify and evaluate the effect of different light stimulation paradigms on the Ca^2+^ dynamics of single cells expressing three existing ChR2 variants i.e. wild type, ChETA, and ChRET/TC. Secondly, to identify key features necessary for the development of prospective ChR2 constructs, we perform a global sensitivity analysis of different parameters of the single cell model to Ca^2+^ spiking rate and basal levels. Thirdly, through the incorporation of gap junctions allowing diffusion of IP_3_ and Ca^2+^, we analyze the effect of local light stimulation on the global Ca^2+^ response in a network of astrocytes homogeneously expressing ChR2. Lastly, we investigate the effect of varying degrees of heterogeneity in ChR2 expression on network-wide astrocytic Ca^2+^ spiking rate and basal level upon global light stimulation.

## Materials and methods

### The biophysical model

In this study, we present a novel biophysical model of optogenetically-modified astrocytes. The model is composed of a combination of the previously published *stochastic astrocyte model* [58, 59] and a *4-state model for ChR2* taken from Stefanescu *et al* [60] and Williams *et al* [61]. The stochastic IP_3_R model is adapted from the Li-Rinzel simplification of the De Young-Keizer model [62–64]. The 4-state ChR2 model assumes the existence of the channel in two closed states (c_1_, c_2_) and two open states (o_1_, o_2_).

**Figure 1.**
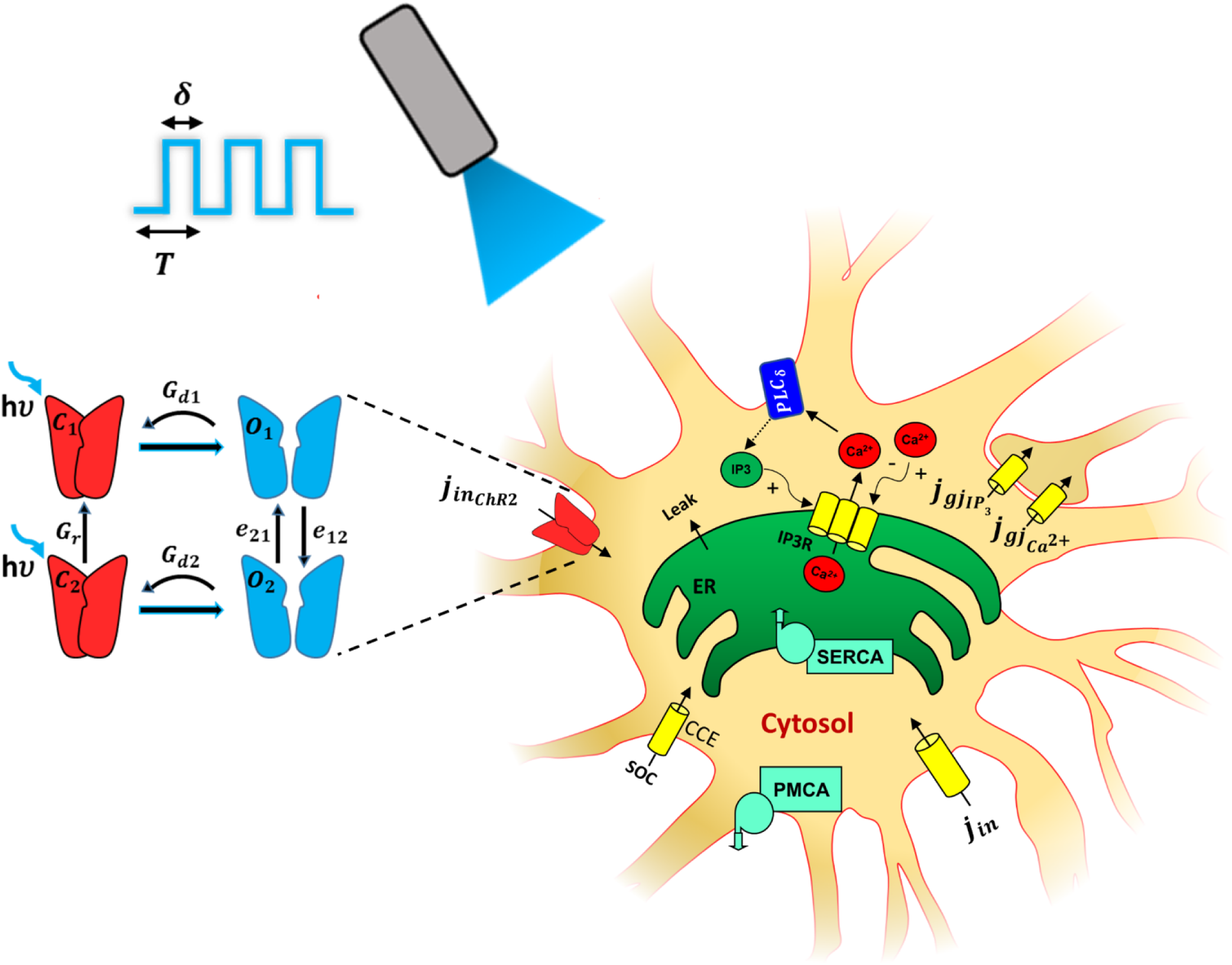
Schematic of the biophysical model of a ChR2 - expressing astrocyte. Inset: The 4-state model of Channelrhodopsin 2 (ChR2) – closed states (c_1_ and c_2_) in red, open states (o_1_ and o_2_) in blue. The rate constants of transitions between states are depicted in the figure. Blue light (hν: 473nm) opens ChR2, facilitating cationic influx 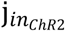, including Ca^2+^, initiating a cascade of Ca^2+^ responses. The light stimulation window is illustrated as a pulse train given by T (pulse period) and δ (pulse width). The model accounts for: 1) Ca^2+^ release from the endoplasmic reticulum (ER) into the cytosol via the IP_3_R clusters, 2) PLCδ mediated production of IP_3,_ 3) capacitative calcium entry phenomenon (CCE) via the store operated calcium channel (SOC), 4) passive leak from the ER to the cytosol, 5) replenishment of ER stores via the SERCA pump, 6) extrusion of Ca^2+^ by PMCA pump (plasma membrane Ca^2+^ ATPase) into the extracellular (EC) space, and 7) passive leak (*j*_*in*_) into the cytosol from the EC. In a network of astrocytes, each cell is connected to its neighboring cells though Ca^2+^ and IP_3_ permeable gap junctions, indicated as 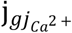 and 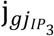, respectively.

Figure 1 illustrates the schematic of the biophysical model of calcium dynamics of ChR2 expressing astrocytes. Cationic influx through ChR2 activation is labeled as 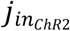. Light stimulation window is modeled as the commonly used pulse train (θ(t)) given by T (pulse period) and δ (pulse width – expressed as percentage of T). Ca^2+^ in the cytosol activates the IP_3_ receptor (IP_3_R) on the endoplasmic reticulum (ER) membrane, leading to an efflux of Ca^2+^ into the cytosol. Cytosolic Ca^2+^ also binds to PLC_δ_ (on the PM) leading to the production of IP_3_ in the cytosol, which also activates IP_3_R clusters. Ca^2+^ release from IP_3_R leads to a further increase of IP_3_R activity, also known as calcium induced calcium release (CICR). Further increase in Ca^2+^ concentration in the cytosol inactivates the release from the ER. Release of Ca^2+^ from the ER leads to capacitative calcium entry (CCE) via the transmembrane store operated calcium (SOC) channel. Uptake of Ca^2+^ via sarcoplasmic reticulum Ca^2+^-ATPase (SERCA) pump results in the replenishment of the ER stores from the cytosol. The PM Ca^2+^ ATPase (PMCA) pump extrudes Ca^2+^ from the cytosol to the extracellular (EC) space.

Our biophysical model for a single astrocyte is composed of nine state variables, i.e. free cytosolic calcium concentration – [Ca^2+^]_c_, inositol triphosphate concentration – [IP_3_], the fraction of open inactivation IP_3_R gates –h, total free Ca^2+^ concentration – c_o_, fraction of ChR2 in its closed and open states – c_1_, c_2_, o_1_,o_2_, and a variable capturing temporal kinetics of conformational changes in ChR2 – s. Additive Weiner processes (σ’s), which capture the stochasticity in astrocytes and ChR2 dynamics, are added as diffusion terms. A network of homogeneous/heterogeneous astrocytes was modeled by incorporation of gap junctions, *J*_*gj*_, between the cells where the diffusion of IP_3_ and Ca^2+^ were accounted for (network dynamics, Table 1). Quantification of spiking rate and Ca^2+^ basal levels were performed pre, during and post stimulus. The equations for astrocyte *‘i’* in the network can be summarized by the following stochastic state-space equation:

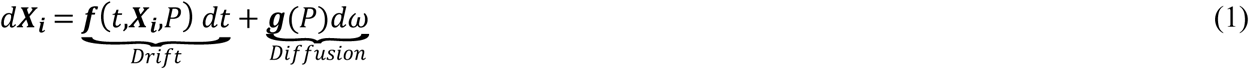

where

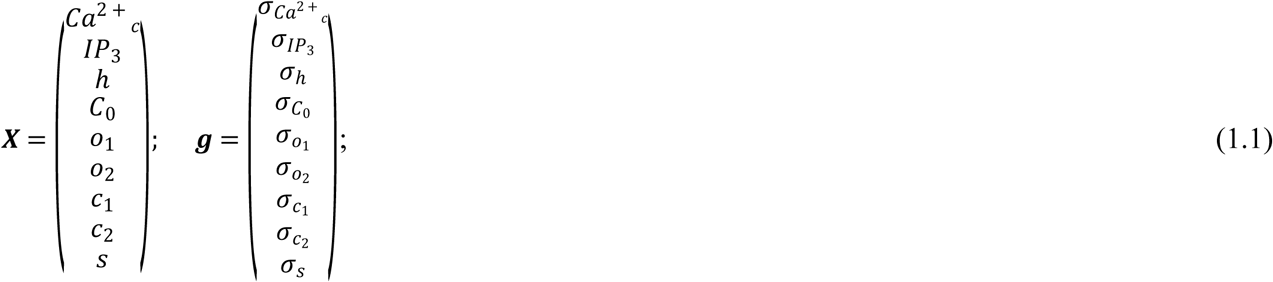

P denotes the parameters of the model, summarized in Table 1, and components of the ***f*** vector will be described in detail. We have previously estimated the variance of the Weiner processes for IP_3_, Ca_c_, h and c_o_, using the local linearization (LL) filter [59, 65] (Weiner processes, Table 1). Potential stochasticity in ChR2 dynamics is included in the model using constant Weiner processes and will be explored in later sections.

**Table 1.**
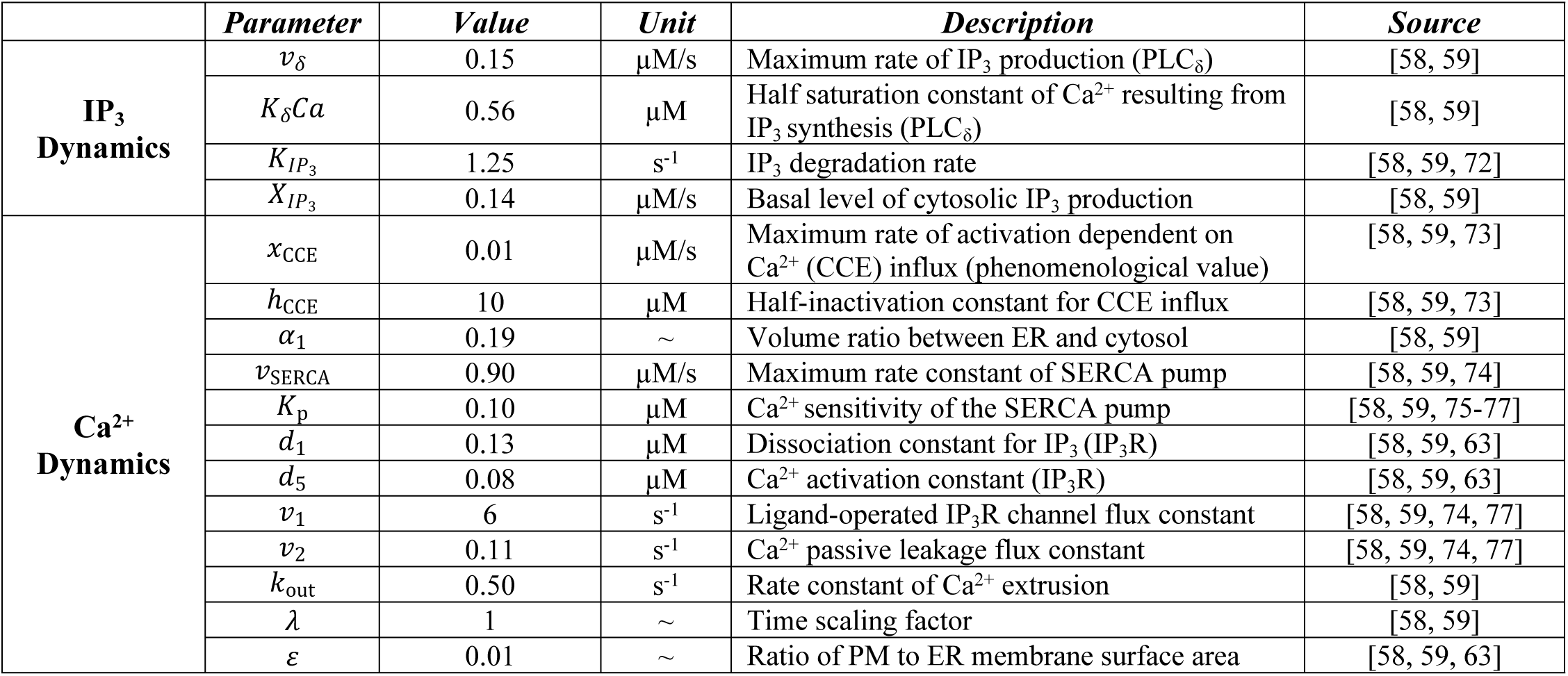
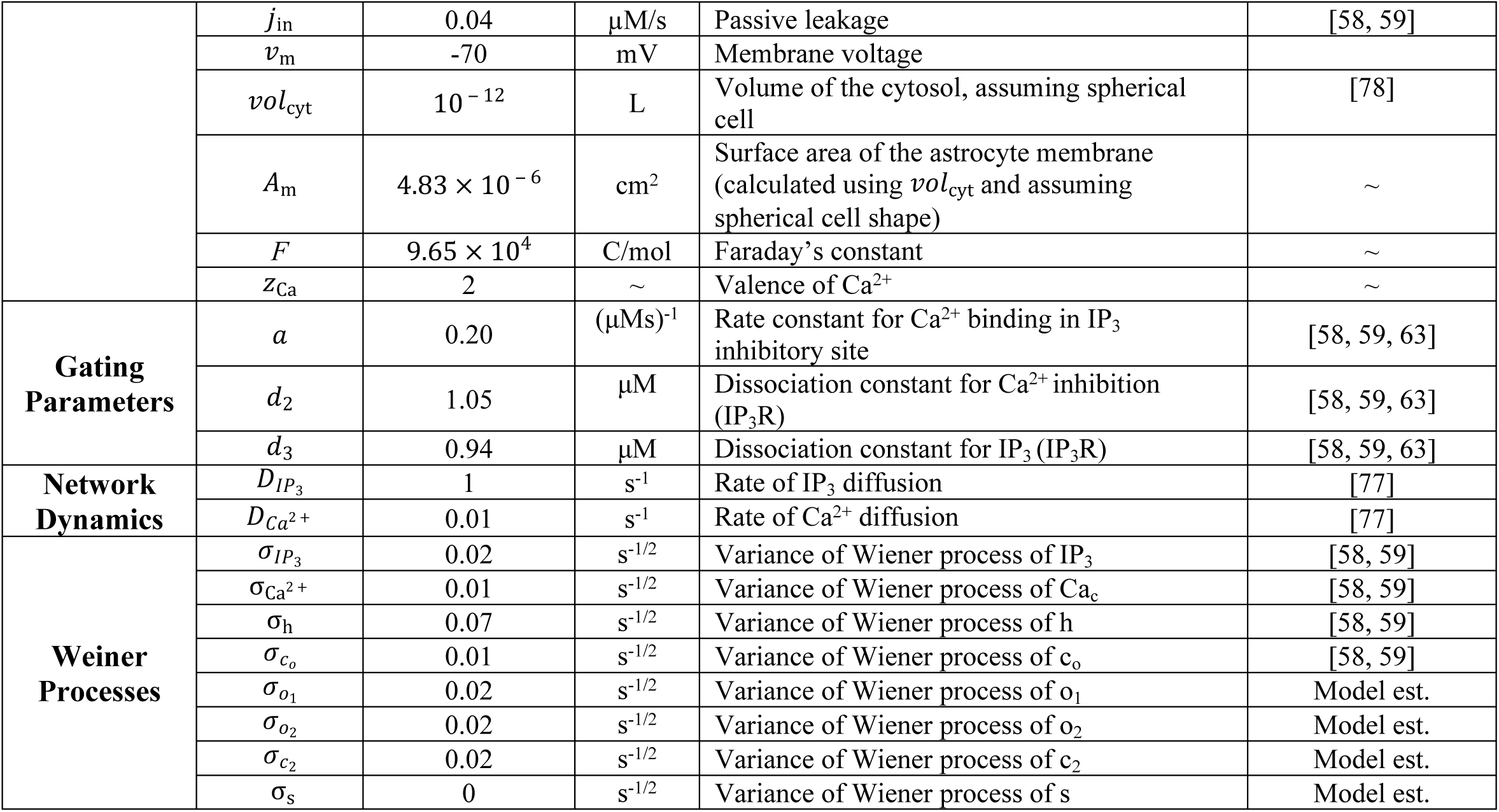
Astrocyte Model Parameters

The dynamics of free cytosolic calcium concentration is given by

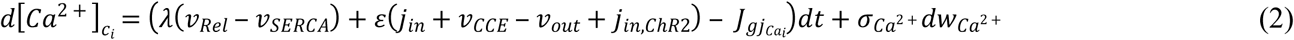

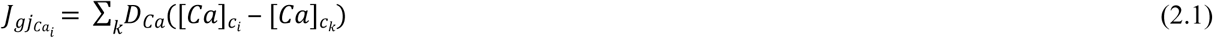

Where 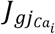 is the gap junctional flux of Ca^2+^ flowing from astrocyte ‘i’ to its neighboring astrocytes (indicated by index k). The efflux of Ca^2+^ from the ER to the cytosol via the IP_3_R is described by

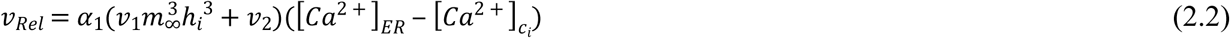

where calcium in the ER is given by

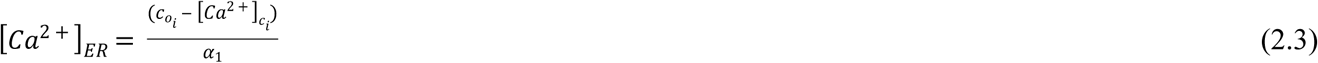

The steady state profile of the open activation IP_3_R gates is

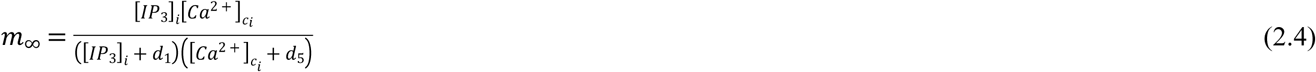

A hill-type kinetic model describing the SERCA pumping is given by

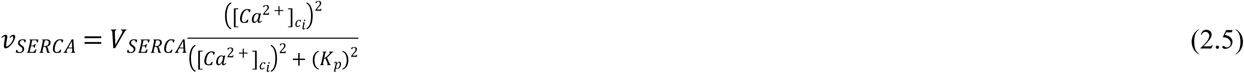

The CCE effect is described as a phenomenological model using the following equation

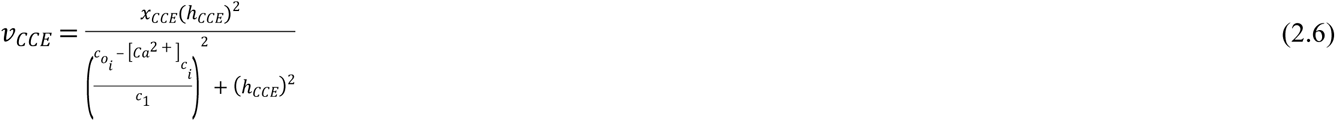

Ca^2+^ extrusion across the PM via PMCA is given by

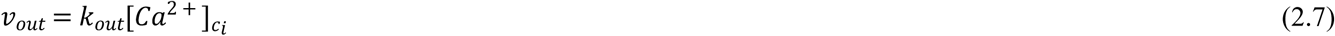

IP_3_ changes in astrocytes mediated by PLC_δ1_ and intercellular diffusion is described as:

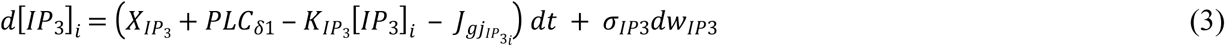

where 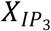 denotes the basal level of IP_3_ production (µM/s) from fluctuations in the action of receptor-agonists over G-protein-coupled receptors, and 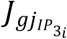 is the gap junctional flux of IP_3_ flowing from astrocyte ‘i’ to its neighboring astrocytes, defined as:

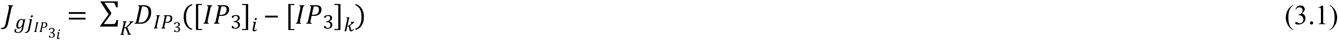

PLC_δ1_ activity is described as the Hill’s kinetic model as

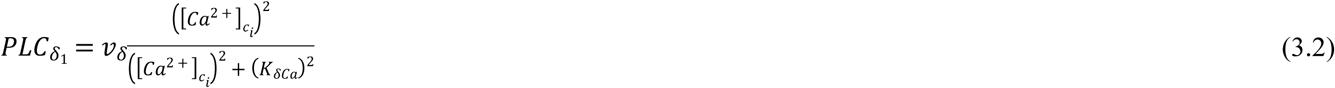

Dynamics of the fraction of open inactivation IP_3_R inactivation gates is given by

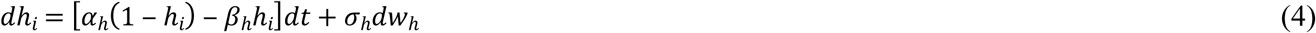

where the opening (*α*_*h*_) and closing rates (*β*_*h*_) rates are

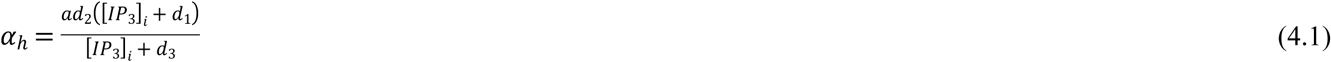

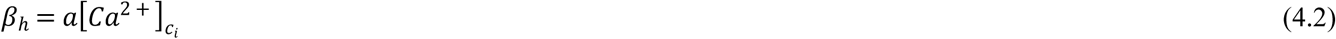

The total free [Ca^2+^] in the cell ([Ca^2+^]_c_ + [Ca^2+^]_ER_) is modeled as

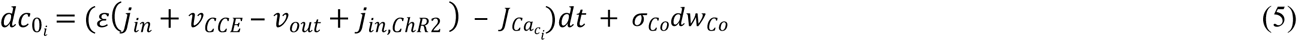

The open and closed gating dynamics of ChR2 are given by equations 5-8, as

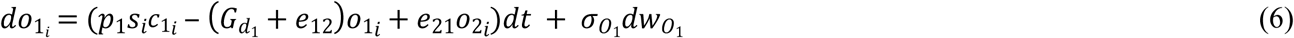

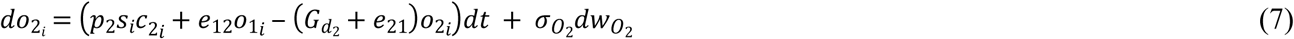

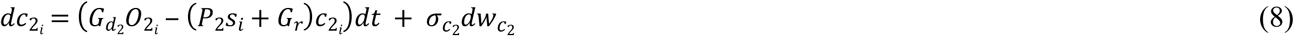

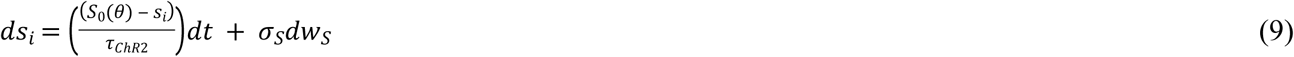

where *θ*(*t*) describes the laser stimulus paradigm as a pulse train, and:

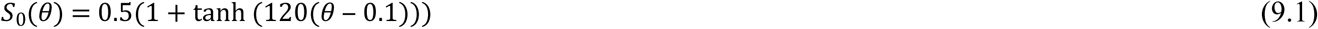

The existence of ChR2 in various states should satisfy the following algebraic condition:

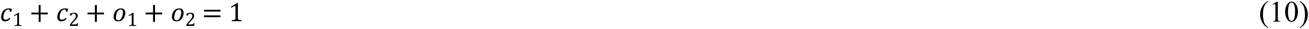

The current generated by cationic influx through ChR2 is given by

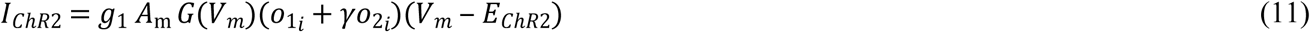

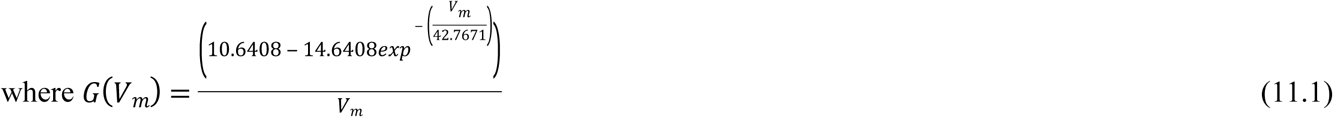

The resultant flux through ChR2 is

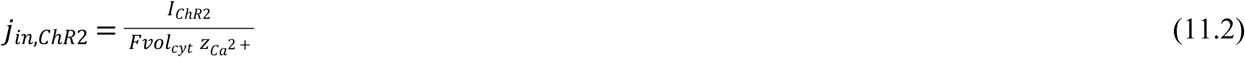

The diffusion term in equation 1 implies solving of the state-space system as an integrated model. In a deterministic system, due to lack of feedback from Ca^2+^ dynamics into that of ChR2, the dynamics of ChR2 can be solved independently. The model was implemented in MATLAB 2018a (Mathworks Inc.) and was numerically solved using the LL method [59] with an integration step size of *Δ*t = 0.1 ms. A listing of all parameters and their descriptions can be found in Tables 1 and 2.

**Table 2.**
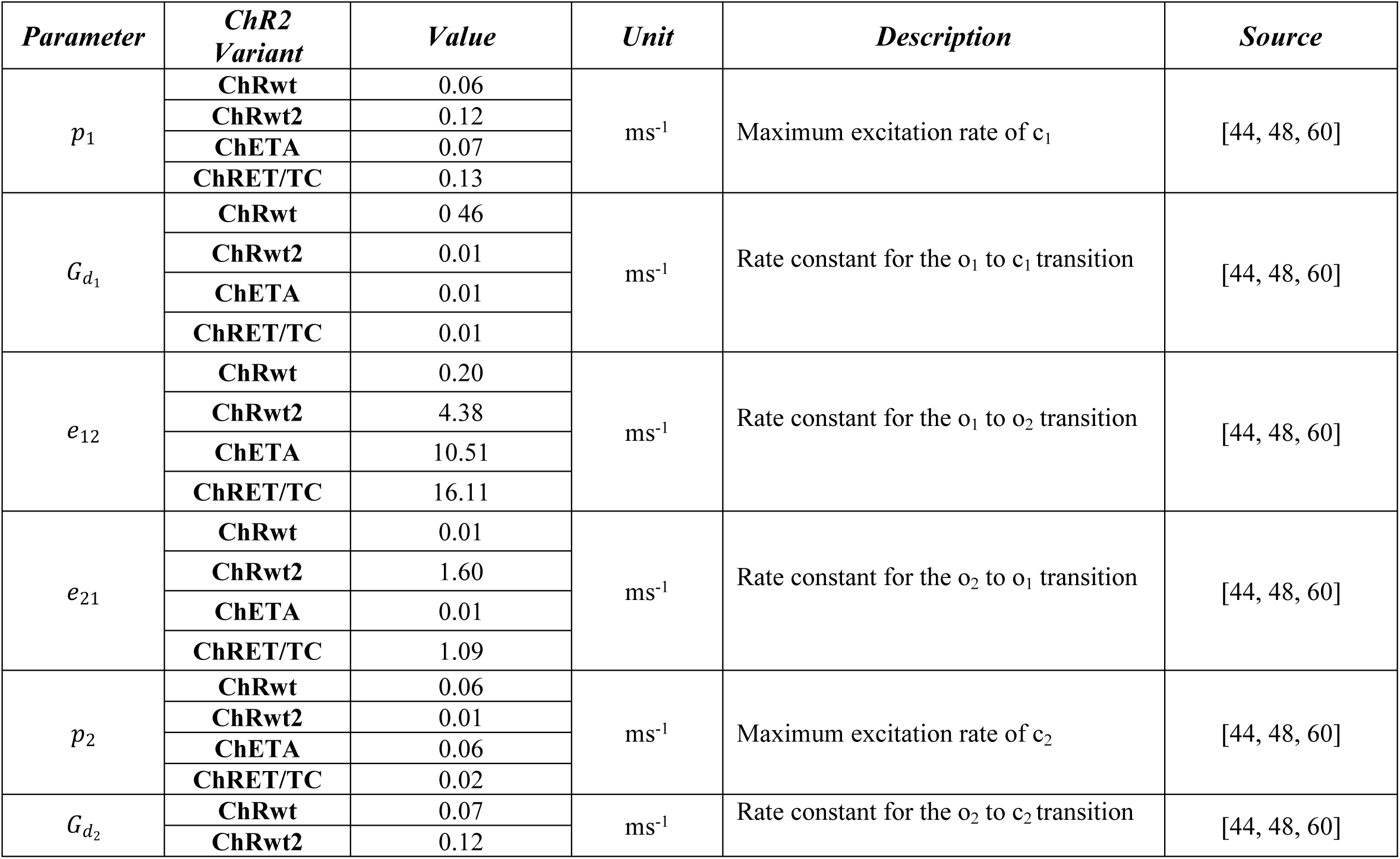
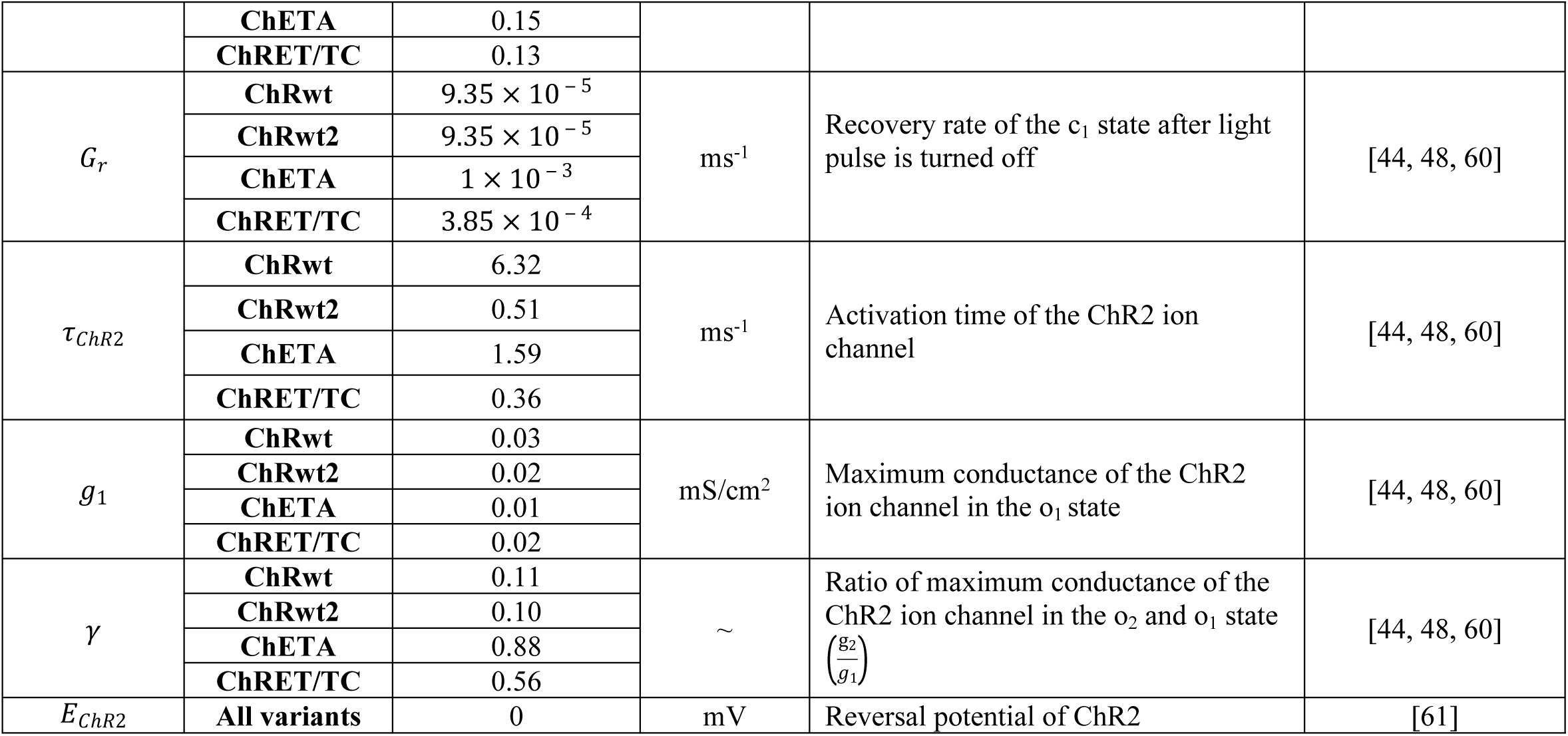
ChR2 4-State Model Parameters

### Light stimulation paradigm

In all simulations performed in this study, laser stimulus was modeled as a square wave pulse train with period T, pulse width δ (expressed as a percentage of T), and unit pulse amplitude. This paradigm is employed to evaluate the effect of light on astrocytic activity, in both individual and a network of gap junction connected astrocytes.

### Sensitivity Analysis

A global sensitivity analysis was performed to assess the sensitivity of SCOs to stochastic noise, without light stimulation. The Latin hypercube sampling (LHS) method with uniform distribution was used to select parameter sets for testing and solving the system [66, 67]. Variance of each of the Weiner processes was varied between a lower and an upper bound (state variable variances, Table 3), and the partial rank correlation coefficient (PRCC) analysis was performed. 95% confidence interval was chosen for statistical significance. A similar global sensitivity analysis was performed to quantify the sensitivity of the Ca^2+^ response to the parameters of ChR2, during light stimulation. Parameter sets accommodating for ranges across parameters were chosen by the LHS method with uniform distribution and the PRCCs were computed with respect to the spiking rate and Ca^2+^ basal level in the astrocyte.

**Table 3.** Global Sensitivity Analysis Ranges for each State Variable Variance and ChR2 Parameter

|  | *Parameter* | *Range* | *Unit* |
| --- | --- | --- | --- |
| **State Variable Variances** | $\sigma_{IP_3}$ | 0 – 0.10 | $\text{s}^{-1/2}$ |
| | $\sigma_{Ca^{2+}}$ | 0 – 0.10 | $\text{s}^{-1/2}$ |
| | $\sigma_h$ | 0 – 0.10 | $\text{s}^{-1/2}$ |
| | $\sigma_{c_o}$ | 0 – 0.10 | $\text{s}^{-1/2}$ |
| | $\sigma_{o_1}$ | 0 – 0.10 | $\text{s}^{-1/2}$ |
| | $\sigma_{o_2}$ | 0 – 0.10 | $\text{s}^{-1/2}$ |
| | $\sigma_{c_2}$ | 0 – 0.10 | $\text{s}^{-1/2}$ |
| **ChR2 Parameters** | $p_1$ | $51.28 - 1.50 \times 10^2$ | $\text{ms}^{-1}$ |
| | $G_{d_1}$ | $8.16 - 5.47 \times 10^2$ | $\text{ms}^{-1}$ |
| | $e_{12}$ | $163.52 - 1.93 \times 10^4$ | $\text{ms}^{-1}$ |
| | $e_{21}$ | $4.00 - 1.31 \times 10^3$ | $\text{ms}^{-1}$ |
| | $p_2$ | $14.08 - 7.70 \times 10^1$ | $\text{ms}^{-1}$ |
| | $G_{d_2}$ | $56.32 - 1.81 \times 10^2$ | $\text{ms}^{-1}$ |
| | $G_r$ | 0.075 – 1.00 | $\text{ms}^{-1}$ |
| | $\tau_{ChR2}$ | 0.000 – 0.0003 | $\text{ms}^{-1}$ |
| | $\gamma$ | 0.000 – 0.011 | ∼ |
| | $g_1$ | 0.0001 – 0.091 | $\text{mS}/\text{cm}^2$ |

## Results

### Response of ChR2 variants to light stimulation

Figure 2 shows a representative simulation of the response to light stimulation of single astrocytes expressing four ChR2 variants - wt1, wt2, ChETA, and ChRET/TC (refer to Table 2 for their gating parameters and conductance). The light stimulation paradigm employed had T = 1s, δ = 20% with a unit pulse amplitude. Within the 20-minute window of simulation, the laser stimulus was applied to the astrocyte from 4-12 minutes. Upon light stimulation (panels A-D), the dynamic system shows increases in [IP_3_], [Ca^2+^] and [c_o_] across ChR2 variants in the order of wt1<wt2<ChETA<ChRET/TC, while the IP_3_R gating variable (h) shows a decrease in the order of wt1>wt2>ChETA> ChRET/TC. In addition, as compared to the pre and the post light stimulus phases, there is an increase in basal Ca^2+^ levels during light stimulation. The ChR2 gating dynamics (panels E-H) indicate that before light stimulation there is a maximum probability of existence of the channel in the c_1_ state. However, upon stimulation, different ChR2 variants show that the dynamic system proceeds to the other states (o_1_, o_2_ and c_2_), with them showing differing gating dynamics.

**Figure 2.**
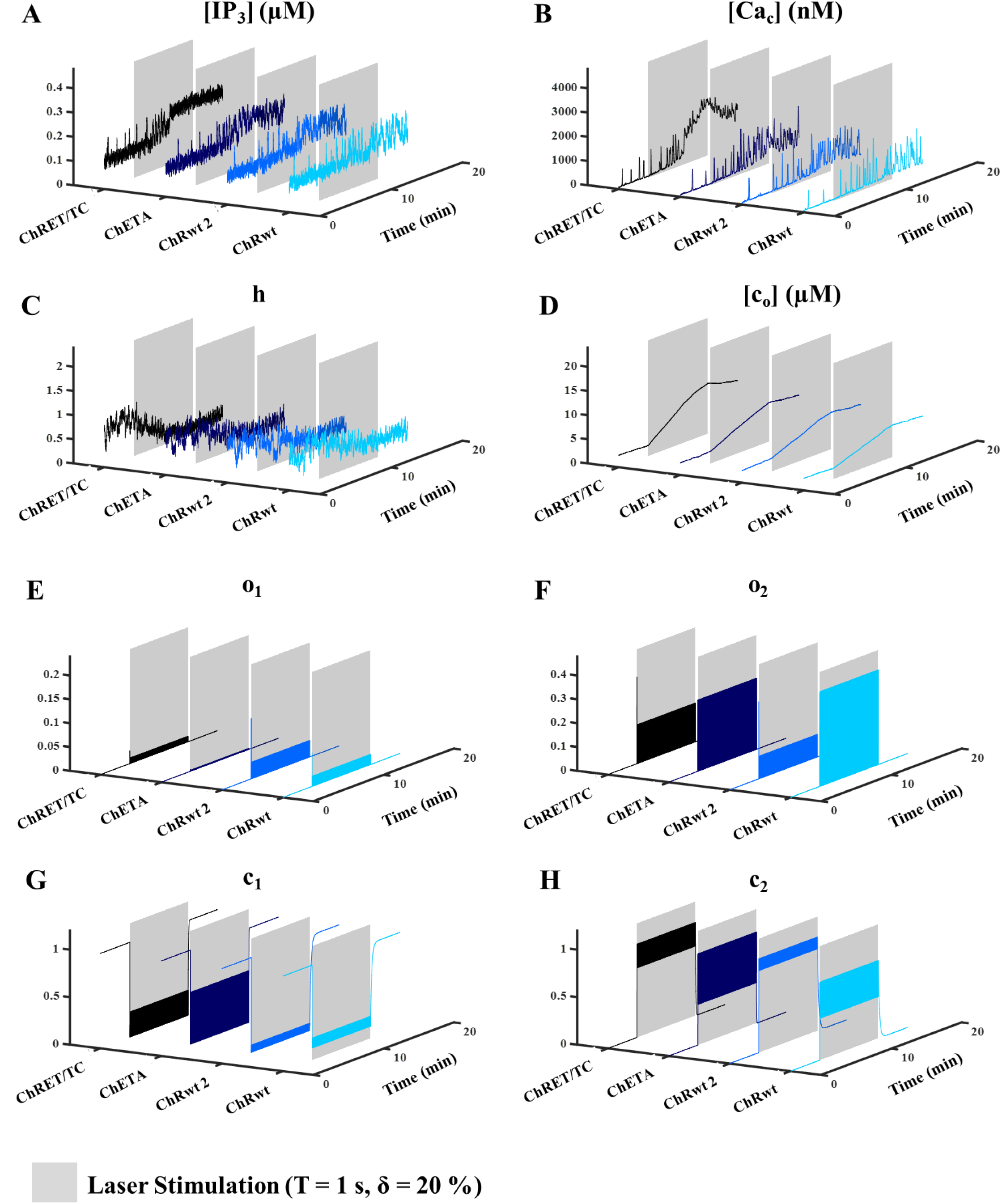
Response of ChR2 variants (wild type 1 (wt1), wild type 2 (wt2), ChETA and ChRET/TC) to light stimulation. The stimulation paradigm (from 4 to 12 minutes, gray shaded region) is a pulse train with the duration (T) = 1 s, pulse width (δ) = 20% (0.2 s), and a unit pulse amplitude. **(A-D)** Representative traces of IP_3_ level ([IP_3_]), cytosolic calcium ([Ca_c_]), inactivation IP_3_R gating variable (h), and total calcium concentration ([c_o_]) for an astrocyte expressing various ChR2 variants upon laser light stimulation are illustrated. Results show high sensitivity of all variables to light stimulation. In particular, increase in [IP_3_], [Ca_c_], and [co], and a decrease in h for all variants during light stimulation is observed. [Ca_c_] traces for the ChRET/TC variant during stimulation shows an elevation of calcium beyond physiological levels, and an apparent reduction in the calcium spiking activity compared to other variants shown in panel B. **(E-H)** Representative traces of the open (o_1_, o_2_) and closed states of ChR2 (c_1_, c_2_) are plotted with respect to time (min), for an astrocyte expressing different variants of ChR2. During pre and post light stimulation phases, all variants have the tendency to stay in the c_1_ state. During the period of stimulus, however, variants show varying degrees of existence in all open and closed states of ChR2.

### Response of a ChETA-expressing astrocyte to various light stimulation paradigms

Figure 3 shows the effect of stimulation paradigms on the mean spiking rate and the steady state Ca^2+^ basal level in astrocytes expressing ChETA (for other ChR2 variants refer to Figures S1-3). Laser parameters (Figure 1) - T and δ were varied between 1-5 seconds and 0-100% of T, respectively. Figure 3A shows a histogram of all Ca^2+^ spikes pre and during light stimulation. To exclude minor irrelevant Ca^2+^ fluctuations, a cutoff prominence (dashed line – 350 nM) was chosen for spiking rate calculations. The cutoff concentration was chosen based on the bimodal distribution of the spiking rate histogram, ensuring that only spikes in the larger mode were chosen and those related to the 1/f noise were excluded from the analysis.

**Figure 3.**
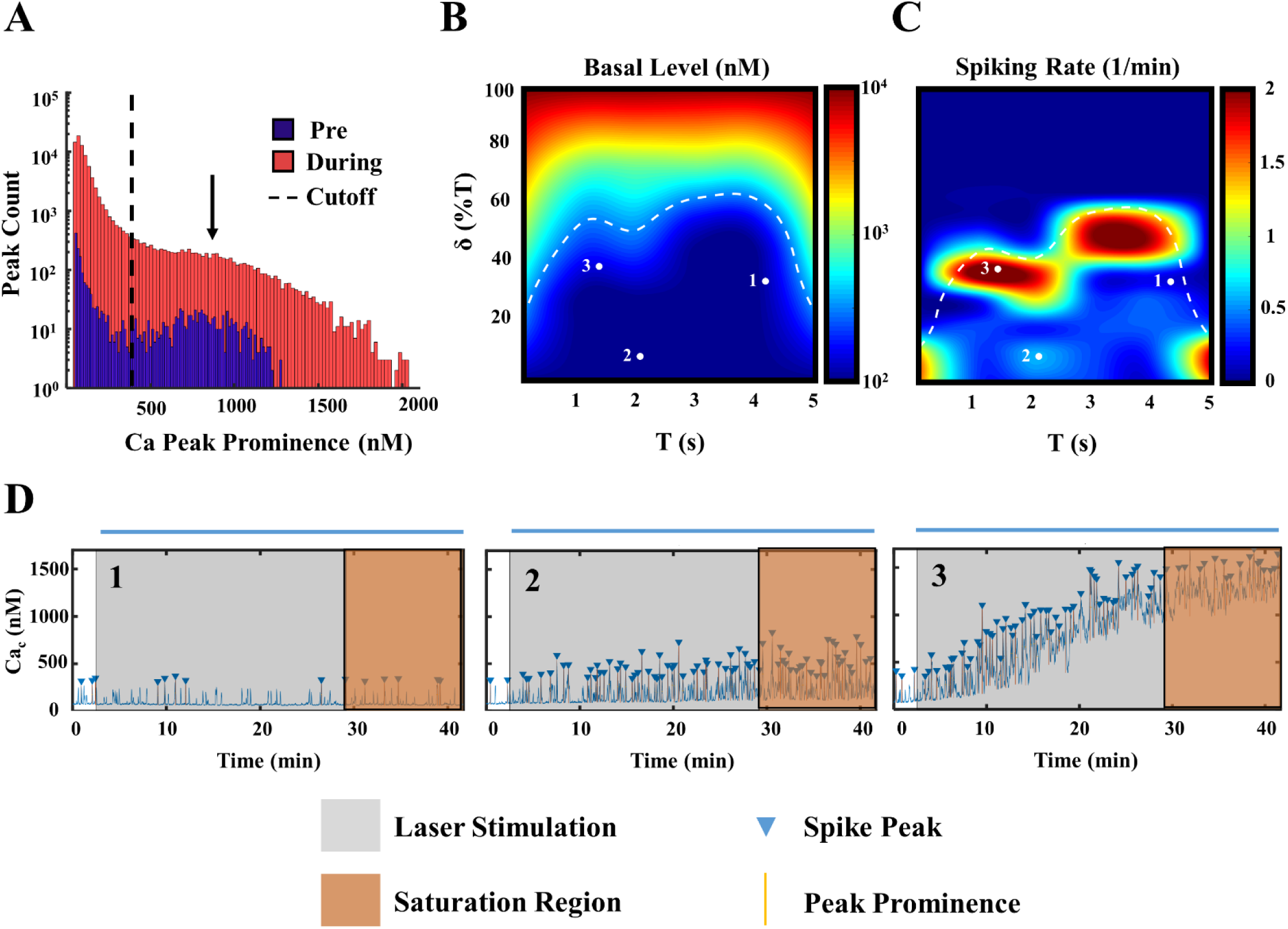
Response of a ChETA - expressing astrocyte to various light stimulation paradigms. **A.** Histogram depicting the peak count in the Ca^2+^ trace of the astrocyte (log scale) with respect to Ca^2+^ peak prominence upon laser light stimulation. Light stimulation parameters – T was varied between 1-5 s; δ between 0-100% of T; unit pulse amplitude. The histogram was generated for the pre-stimulus phase (blue) and during stimulus phase (red). The cutoff prominence was set to 350 nM, in accordance with the observed bimodal distribution of Ca^2+^ spikes (dashed line), and to assure that 1/f noise related Ca^2+^ spikes are not included in the analysis. **B.** The T-δ heat map of the Ca^2+^ basal level for various combinatorial windows of T and δ, expressed in the log scale. Specific regions in the physiological levels of Ca^2+^ basal level (indicated by the white dashed trace) are numbered and used for further plotting and analysis. **C.** The T-δ heat map indicating spiking rate in the astrocyte for various combinatorial windows of T and δ, above the cutoff prominence chosen in (**A**). White dashed trace delimits the physiological basal levels; as defined in (**B**). **D.** Representative Ca^2+^ signaling traces of points 1, 2 and 3, from (**B**) and (**C**). Light stimulation was started at 50 s until the end of the simulation (blue bar). Mean Ca^2+^ spiking rate across trials was calculated once the Ca^2+^ signal trace reached a steady profile (in orange).

For each combination of T and δ, 10 trials were performed for 40 minutes, with the light stimulation starting at 50 seconds until the end of the simulation (indicated by the grey window and the blue bar). Once the Ca^2+^ baseline reached a steady profile (indicated by the orange region), the mean spiking rate across trials and mean basal levels were calculated for each stimulation paradigm. The T-δ heat (color) maps, useful to determine optimal Ca^2+^ signaling behavior in astrocytes exposed to a variety of T and δ combinations, are shown in Figure 3B and C for Ca^2+^ baseline and Ca^2+^ spiking, respectively. Results indicate that in the physiologically acceptable ranges chosen for T and δ (Ca^2+^ basal levels higher than reported physiological values are separated by the dashed white trace in panel B), there are regions of increased astrocytic Ca^2+^ spiking activity (red regions in Figure 3C). Three representative traces from regions with low, intermediate and high astrocytic Ca^2+^ spiking activity with physiological Ca^2+^ basal levels are depicted in Figure 3D.

### Sensitivity of the astrocytic Ca^2+^ response to system state variables and ChR2 parameters

In Figure 4, a global sensitivity analysis was performed to evaluate the Ca^2+^ response of astrocytes (i.e. the spiking rate and steady basal level) to variations in the stochastic noise variances of the state variables (without light stimulus) as well as to variations in the parameters of ChR2 (during light stimulation). To generate Figure 4A, simulations were performed for 10 minutes, for 10 trials without light stimulation. The range of the variances (σ’s) were chosen between 0 and 0.10 (state variable variances, Table 3). The LHS method with uniform distribution was used to choose 500 parameter sets for simulations. The PRCC of the variances of the Wiener processes with respect to the mean Ca^2+^ spiking rates, along with the corresponding p-values were computed. As seen, SCOs are highly sensitive (indicated by *) to the variances in the order of 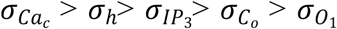; however, the contributions of 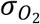 and 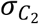 were not significant.

**Figure 4.**
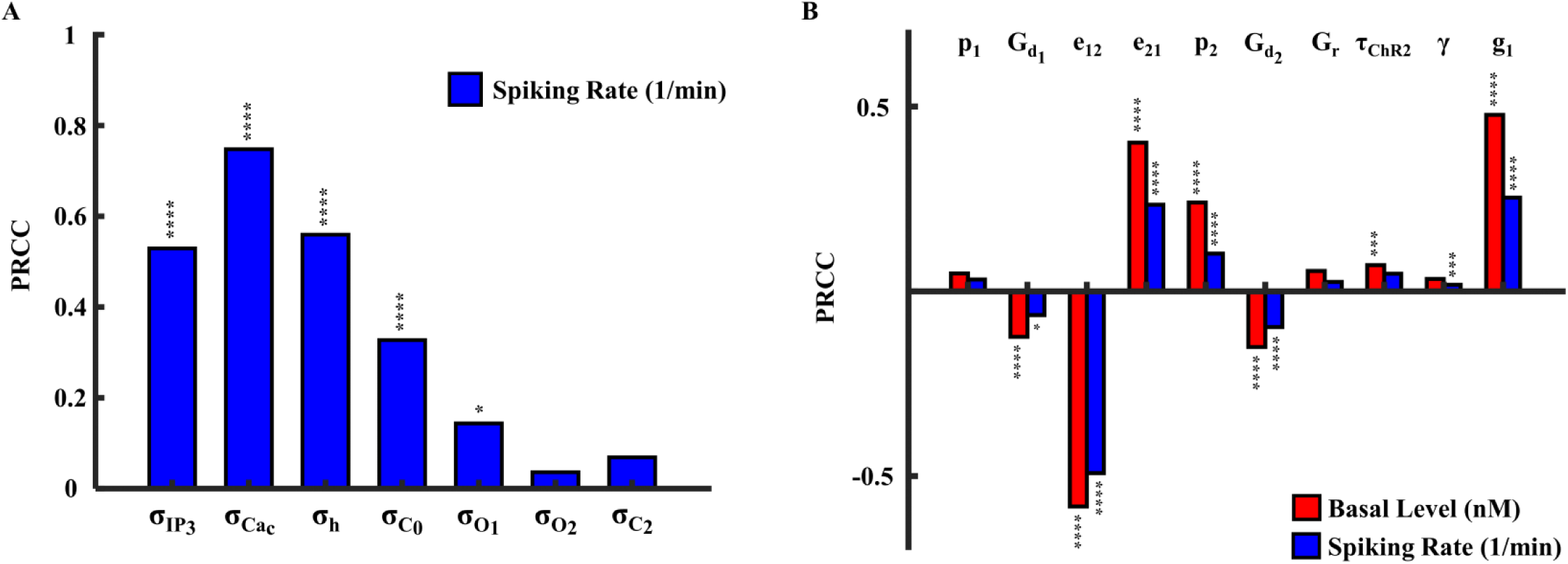
Sensitivity of the astrocytic Ca^2+^ response to the state variable variances and ChR2 parameters. **A.** Global sensitivity analysis results depicting sensitivity of astrocyte Ca^2+^ response to stochastic noises, without light stimulation. Partial rank correlation coefficients (PRCCs) with respect to the Weiner processes of the state variables are plotted. 500 parameter sets chosen by the Latin hypercube sampling (LHS) method with uniform distribution. * depicts significance levels. Spiking rate *p*-values: 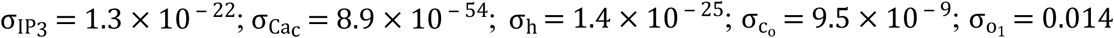.**B.** Plot of the PRCCs for each parameter of ChR2 during light stimulation (T= 4.5 s, δ = 1.35 s (30% of T), light stimulation started at 50 s and continued for the duration of the simulation, (total simulation time = 40 min, 10 trials) with respect to the basal level (nM) and spiking rate (1/min); peak prominence = 350 nM. 1000 parameter sets were chosen using the LHS sampling method with uniform distribution. Spiking rate p-values: 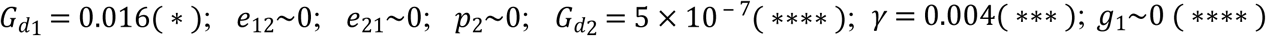. Basal level p-values: 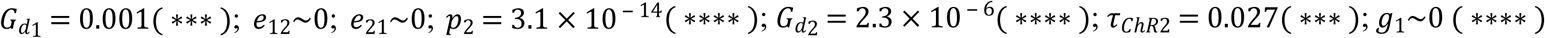.

Figure 4B shows the sensitivity of Ca^2+^ activity to parameters of ChR2 during light stimulation with the paradigm shown in Figure 3D, trace 1 (T = 4.5s and δ = 30%). Similar to the analysis in Figure 3A, the cutoff prominence of the peaks counted was set to 350 nM to exclude 1/f noise related Ca^2+^ spikes. The range of each parameter was chosen such that the four ChR2 variants were encompassed in it (ChR2 parameters, Table 3). 1000 parameter sets were chosen using the LHS method with uniform distribution. For each parameter set, 10 trial simulations were performed, each with a duration of 40 minutes, and the respective Ca^2+^ spiking rate and steady basal levels were calculated. Light stimulation was initiated at 50 seconds and continued for the duration of simulation. The PRCCs were computed and plotted for each of the ChR2 parameters evaluated in this study (Figure 4B). The results indicate that the parameters e_12_, e_21_ and g_1_ are statistically significant at a 95% confidence interval, for both, the Ca^2+^ spiking rate and basal level. While e_12_, G_d1_ and G_d2_ are negatively correlated to the Ca^2+^ response with respect to both the spiking rate and basal level, e_21,_ p_2_ and g_1_ are positively correlated. Parameters *τ*_*ChR*2_ and *γ* are statistically significant and positively correlated to the Ca^2+^ response with respect to the basal level and spiking rate, respectively.

### Network-wide response of homogenously ChR2-expressing astrocytes to light stimulation

Figure 5 shows the effect of light stimulation on a network of 10×10 astrocytes homogenously expressing the ChR2 variant - ChETA. Astrocytes are connected to each other in all orientations, i.e., horizontal, vertical and diagonal directions. Light stimulation was performed with T = 2s and δ = 15% and between 12 – 25 minutes (indicated by the grey shaded region, blue bar). The spiking rate in response to the light stimulation was computed as a network-wide behavior throughout the total period. Figure 5A shows a histogram of all Ca^2+^ spikes, pre and during light stimulation, revealing also bimodal distributions. Similar to previous simulations, to exclude irrelevant Ca^2+^ fluctuations due to 1/f noise, a cutoff prominence (dashed line – 350 nM) was chosen for spiking rate calculations. A representative trace of the cytosolic Ca^2+^ is shown in Figure 5B, with the properties of the trace used for quantification, i.e. peak prominence and peaks detected. The grey shaded region represents the time period during which laser stimulation was performed. Network-wide responses to light stimulation, quantified by the mean spiking rate for each cell are shown as heat maps, which were calculated for the pre, during and post light stimulation phases (Figure 5C). The network Ca^2+^ baselines at three specific time instances within the pre, during and post stimulus phases, are shown in Figure 5D (for the full video, refer to supplementary video S1). The spatial arrangement of astrocytes and the subnetwork of astrocytes being stimulated are shown in Figure 5E. Astrocytes along the diagonal (red) are labeled 1 through 6. The Ca^2+^ activity traces of each cell along the diagonal are shown in Figure 5F. Results indicate that there are SCOs across the network, prior to light stimulation, while during stimulation, the astrocytic Ca^2+^ spiking rate is maximum in and around the area of light stimulation. Post light stimulation, the activity is dispersed throughout the network and lasts for long periods of time (Figure 5 B, C and F). Inspection of Ca^2+^ activity traces of various cells across a diagonal with increasing distance from the center of the light stimulation area indicates that there is a decrease in Ca^2+^ spiking rate (Figure 5F). This suggests that there is a propagation of the Ca^2+^ spiking probability resulting in a network-wide effect on Ca^2+^ dynamics.

**Figure 5.**
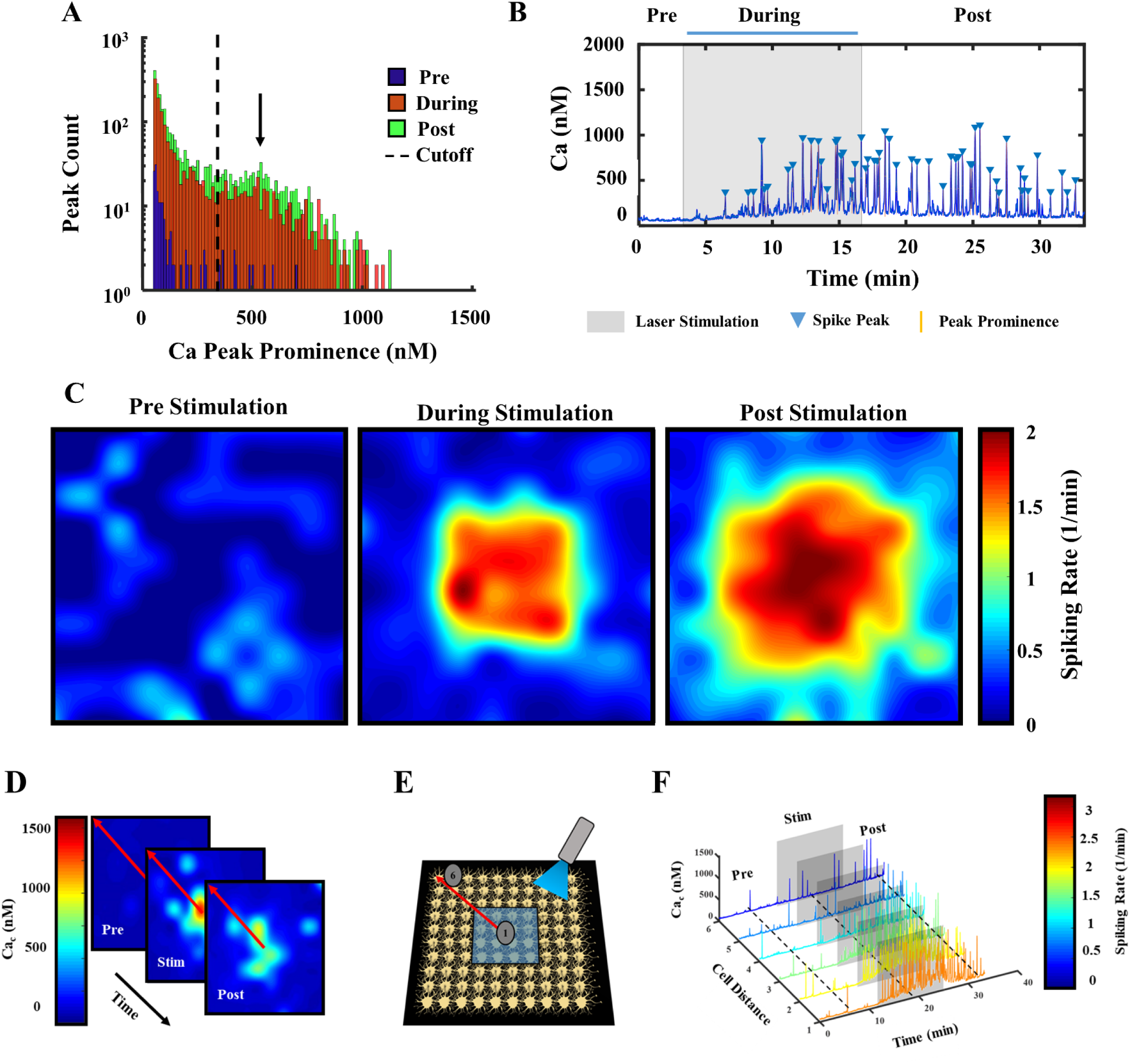
Network-wide behavior of astrocytic Ca^2+^ responses to light stimulation. **A.** Histogram (log scale) depicting the peak count in the Ca^2+^ traces in a 10 x 10 network (100 astrocytes) homogenously expressing ChETA with respect to the Ca^2+^ peak prominence during the pre-stimulus (blue), during stimulus (red) and post stimulus (green) phases. Light stimulation parameters – T = 2 s; δ =15%; unit pulse amplitude. The cutoff peak prominence was set to 350 nM (dashed line) due to the bimodal distribution of Ca^2+^ spikes and assures that 1/f related irrelevant Ca^2+^ spikes are excluded from the analysis. **B.** A representative trace showing the Ca^2+^ signaling profile over time. Light stimulation was performed between 12 – 25 min. (grey shaded region, blue bar). **C.** Heat map indicating the mean Ca^2+^ spiking rate above the cutoff prominence (indicated in (**A**) in the network - pre, during and post light stimulation. **D.** Heat map indicating the Ca^2+^ basal levels in the network – pre, during and post light stimulation. Astrocytes oriented across a diagonal (indicated by red line) were used for further interpretation in (**F**). **E.** Illustration of 4 x 4 subnetwork of astrocytes focally stimulated by blue laser light (indicated by the blue shaded region). 6 astrocytes across the diagonal (in the direction of the red arrow), were used to evaluate the effect of distance from the stimulation on cytosolic Ca^2+^ profiles. **F.** Depiction of Ca^2+^ signal profiles of these 6 astrocytes (in **E**), plotted as a function of time; light stimulation window in grey, color bar represents the mean spiking rate.

### Effect of ChETA-expression heterogeneity on network-wide light stimulation

Figure 6 shows the effect of light stimulation on a network of 5×5 astrocytes with varying degrees of expression of ChETA. The astrocytes are connected in all directions via gap junctions. For each expression level, five random distributions of ChR2 expressing astrocytes were generated, and simulations were performed (15 minutes, 5 trials). Light stimulation was performed from 50 seconds until the end of the simulation. Once Ca^2+^ baseline reached a steady profile, the mean and standard deviation of spiking rates and Ca^2+^ basal levels were computed. Figures 6A (T = 5s, δ = 40%) and 6B (T = 2.5s, δ = 10%) show an increase in Ca^2+^ basal levels and spiking rate in the network, corresponding to increases in ChR2 expression levels. It is to be noted that in the abovementioned light stimulation paradigms, the Ca^2+^ basal levels are within physiological levels (indicated by dashed line). Although Figure 6C (T = 2s, δ = 40%) showed an increase in the Ca^2+^ baseline when the ChR2 expression was increased, there is an overshoot beyond physiological levels at the 80 and 100 % expression levels. Ca^2+^ spiking rate, on the other hand, shows an initial increase until 50% expression level, post which displayed a declining trend.

**Figure 6.**
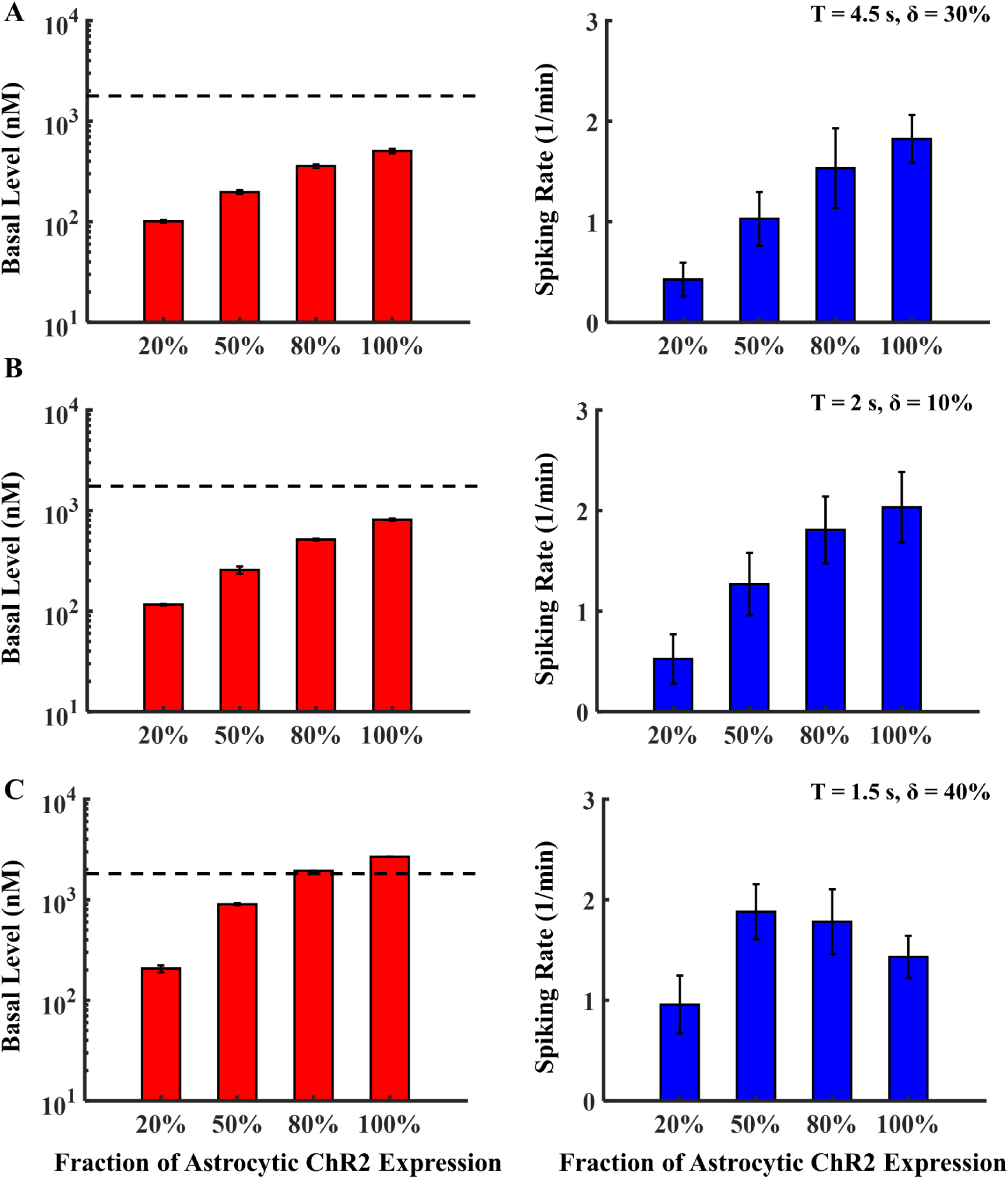
Effect of ChETA expression heterogeneity on network-wide light stimulation. Each bar chart shows the mean network basal level (nM) and spiking rate (1/min) as a function of astrocyte ChR2 expression fraction. Each part corresponds to network-wide stimulation with 1 of 3 different paradigms: **A.** point 1 of Figure 3 (T = 4.5 s, δ = 1.35 s (30% of T), low Ca^2+^ activity), **B.** point 2 of Fig. 3 (T = 2 s, δ = 0.2 s (10% of T), intermediate Ca^2+^ activity), and **C.** point 3 of Figure 3 (T = 1.5 s, δ = 0.6 s (40% of T), high Ca^2+^ activity). In all 3 cases, the stimulation was initiated at 50 s and continued for the duration of the simulation, and the black dashed line marks the maximum physiological basal level of astrocytes.

## Discussion

We developed a novel stochastic model to assess the effect of light stimulation on the Ca^2+^ dynamics in astrocytes expressing the widely used opsin - ChR2. We used three ChR2 variants - wild type, ChETA, and ChRET/TC. The proposed framework can further be adopted for investigating other opsins. Our model accounts for major intracellular calcium signaling pathways as well as light-activated cationic influx through ChR2. We studied light-induced Ca^2+^ responses in both a single astrocyte and a network of homogenously/heterogeneously ChR2 expressing astrocytes. We identified favorable light stimulation paradigms for the abovementioned ChR2 variants which result in maximal spiking rates in astrocytic Ca^2+^ activity within physiological Ca^2+^ basal levels. We also quantified the sensitivity of the model output to changes in the regulation kinetics and in the conductance of ChR2. The model presented in this study provides an insight into stimulation paradigms ideal for controlling astrocytic Ca^2+^ activity and offers geneticists an efficient theoretical framework for the design of new variants.

Results show that calcium dynamics in astrocytes, as seen in experimental studies [51], can be heavily regulated by light-induced activation of ChR2 (Figures 2-3 and S1-3). According to our findings, all ChR2 variants studied in this paper showed similar profiles of activity in response to different laser pulse specifications (Figure 3, S1–3). The profiles displayed common regions of high spiking rate (point 3, Figure 3C), as well as regions with intermediate (point 2, Figure 3C) and low activity (point 1, Figure 3C). Also, with the increase in δ, there is a consistent increase in the basal level observed at any given T. This drastic variability of astrocytic model response to varying stimulation paradigms emphasizes the importance of choosing ‘ideal’ T and δ for desired astrocytic activity in future studies. A wrong selection of these two parameters could prompt these cells to an unhealthy Ca^2+^ signaling regime.

Global sensitivity analysis (Figure 4A) indicates that SCOs are significantly dependent on the stochasticity of IP_3_ dynamics, PM fluxes and the o_1_ state of ChR2. Similar dependencies to IP_3_ receptor activity and membrane fluxes have been shown by us [59] and in a recent study by Ding *et al* [57]. Although the source of stochasticity in ChR2 dynamics is yet to be investigated, we hypothesize that potential protein thermal noise and fluctuations in light intensity due to photon migration dynamics may play a role. Figure 4B indicates that the kinetics of ChR2 significantly affect Ca^2+^ spiking rate and basal level. As a general trend, intuitively, directing ChR2 to the open states (o_1_ and o_2_) from the closed states (c_1_ and c_2_) leads to an increase in astrocytic activity in response to light stimulation. For instance, decreasing 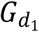 and 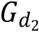 facilitates the existence of ChR2 in the open states as they are negatively correlated to the basal level and spiking rate. Also, increase in p_2_ drives the system to the open state. Similarly, increase in the conductance of ChR2 results in enhanced ionic influx into the cell, thereby elevating both spiking rate and basal levels of calcium. However, less intuitively, increased astrocytic activity occurs when ChR2 exists in the o_1_ state as compared to the o_2_ state. This can be observed, as an increase in e_21_ and decrease in e_12_ led to the existence of ChR2 in o_1_ state (see Figure 1 inset). Collectively, our results suggest that for the light stimulation paradigm used in our global sensitivity analysis, maximal astrocytic activity can be achieved when ChR2 is directed towards the o_1_ state, which can be used for future development of ChR2 variants.

An important aspect in experimental optogenetics is ChR2 expression levels (e.g., transduction efficiency). While incorporating genetic material into the cell, heterogeneity in the degree of expression might occur [68, 69]. Model results suggest that network-wide Ca^2+^ response in astrocytes to light stimulation depends heavily not only on the expanse of stimulation and specification of the paradigm, but also on the degree of heterogeneity (Figures 5, 6, S1–3). We observed the propagation of the probability of Ca^2+^ spiking in response to local light stimulus in a network of homogeneously ChR2 - expressing astrocytes (efficiency of 100%, Figure 5). In heterogeneously ChR2 – expressing astrocytes subjected to a given network-wide stimulus paradigm, differing degrees of heterogeneity resulted in varying degrees of Ca^2+^ spiking and basal levels (Figure 6). The expected increase in the Ca^2+^ spiking rate with the increase in the fraction of ChR2 expression was observed in stimulation paradigms corresponding to points 1 and 2 in Figure 3 (Figure 6A and B). However, notably, due to saturation of astrocytic Ca^2+^ signaling, i.e., elevation of the Ca^2+^ baseline beyond physiological levels, there is a counteracting effect on Ca^2+^ spiking rate when expression is increased (Figure 6C at 80 and 100%). This indicates that design of experiments for stimulation of a network of genetically altered astrocytes cannot be solely based upon observations from single cells, as other factors like ChR2 expression levels and the specific stimulation design (T, δ) play a significant role.

The model presented in this paper aimed at studying the effect of light stimulation on Ca^2+^ dynamics in optogenetically-enabled astrocytes. The model does not include the dynamics of other major ionic species crucial in the function of these cells. Furthermore, membrane electric potential dynamics are not included in the current model, and hence voltage gated calcium channels have not been incorporated. This modeling approach can be applied to other ChR2 variants upon availability of quantified parameters. We sought to provide a minimalistic theoretical framework which can readily be employed by researchers for the investigation of light induced Ca^2+^ responses in astrocytes. Combination of the presented model with more detailed models as in Savtchenko *et al* [70] and Lallouette *et al* [71] where exhaustive geometry and dynamics of various ionic species important for astrocytic Ca^2+^ signaling are accounted for, can enhance our understanding of the intricacies of the behavior of these cells and their response to light.

## Acknowledgments

This study was supported by the National Institutes of Health (1R56NS094784-01A), Wallace Coulter Foundation – BME SEED grant – Florida International University (FIU), and Dissertation Year Fellowship (DYF) from the University Graduate School at FIU (Arash Moshkforoush). The authors would like to sincerely thank Dr. James Schummers, Dr. Nikolaos Tsoukias, Dr. Rita Alevriadou, Mr. Ramkrishnan Krishnan, and Mr. Ricardo Siu for insightful discussions.

## Author contributions

Model development: AM LB CM JS JR. Analysis/discussion of results: CM AM LB JR. Literature review: LB AM CM. Manuscript writing: LB AM CM JR.

## Supplementary Figures -Effect of light stimulation on different ChR2 variants

**Figure S1.**
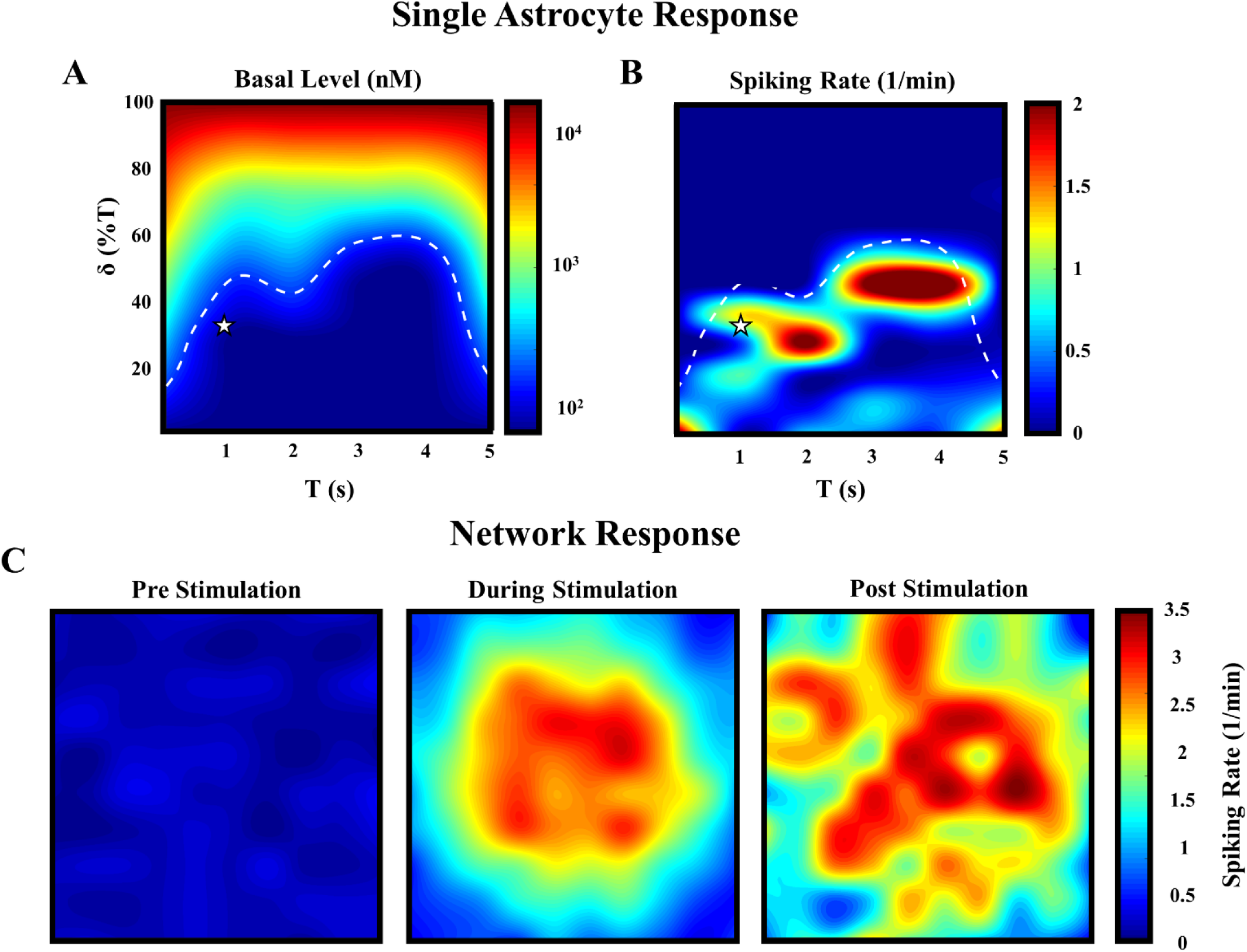
Response of ChRET/TC -expressing astrocytes to light stimulation. **A.** Heat map of the Ca^2+^ steady state basal level for various combinatorial windows of time duration (T) and pulse widths (δ; expressed as a percentage of T), expressed in the log scale. The physiological levels of Ca^2+^ basal level are indicated by the white dashed line. **B.** Heat map indicating spiking rate in the astrocyte for various combinatorial windows of T and δ, above the cutoff prominence chosen in Figure 3A. **C.** Heat maps indicating the mean spiking rate of astrocytes in the network - pre, during and post light stimulation (T = 1 s, δ = 0.3 s (30% of T), the point denoted by star in **A** and **B**.

**Figure S2.**
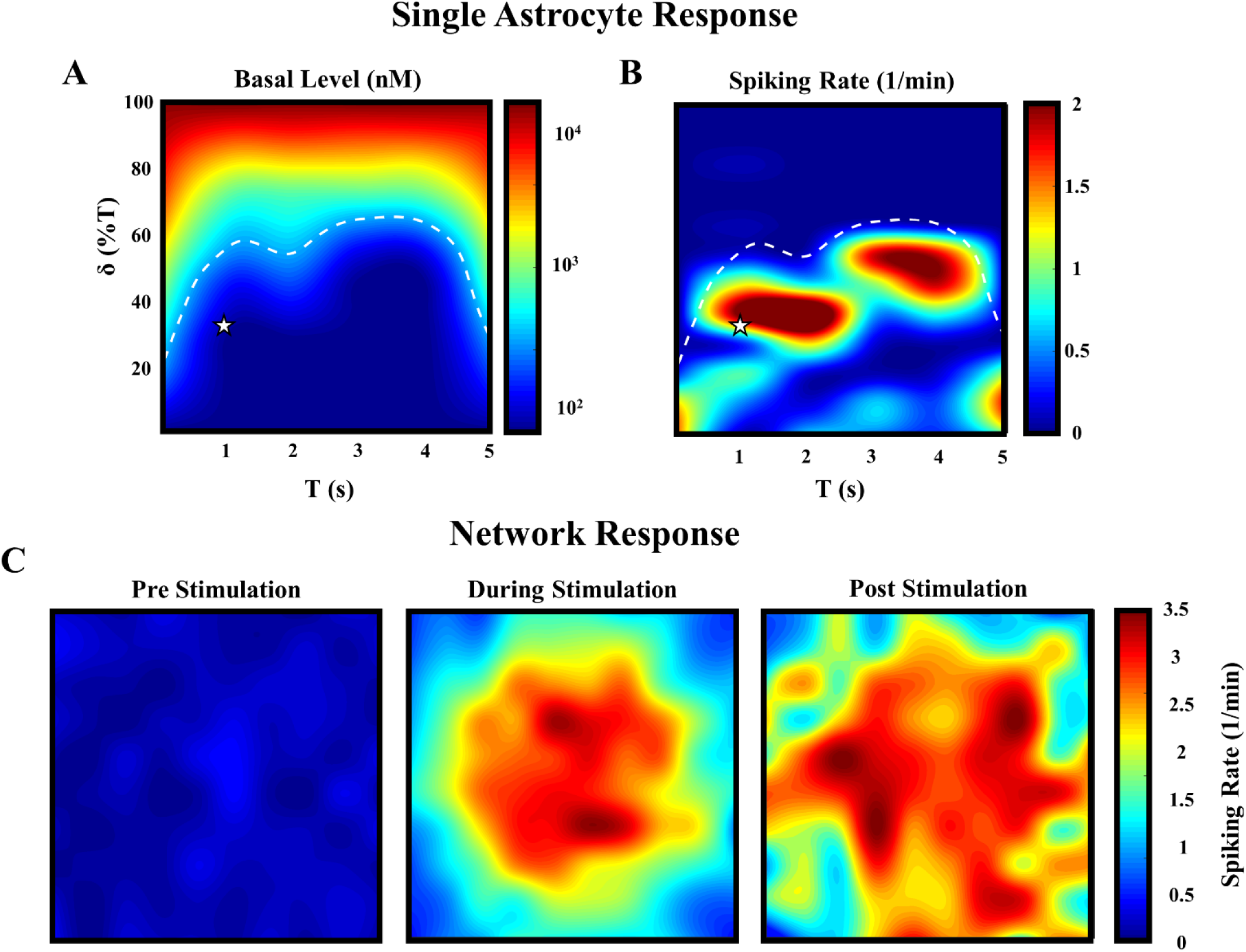
Response of ChRwt1 - expressing astrocytes to light stimulation. **A.** Heat map of the Ca^2+^ steady state basal level for various combinatorial windows of time duration (T) and pulse widths (δ; expressed as a percentage of T), expressed in the log scale. The physiological levels of Ca^2+^ basal level are indicated by the white dashed line. **B.** Heat map indicating spiking rate in the astrocyte for various combinatorial windows of T and δ, above the cutoff prominence chosen in Figure 3A. **C.** Heat maps indicating the mean spiking rate of astrocytes in the network - pre, during and post light stimulation (T = 1 s, δ = 0.3 s (30% of T), the point denoted by star in **A** and **B**.

**Figure S3.**
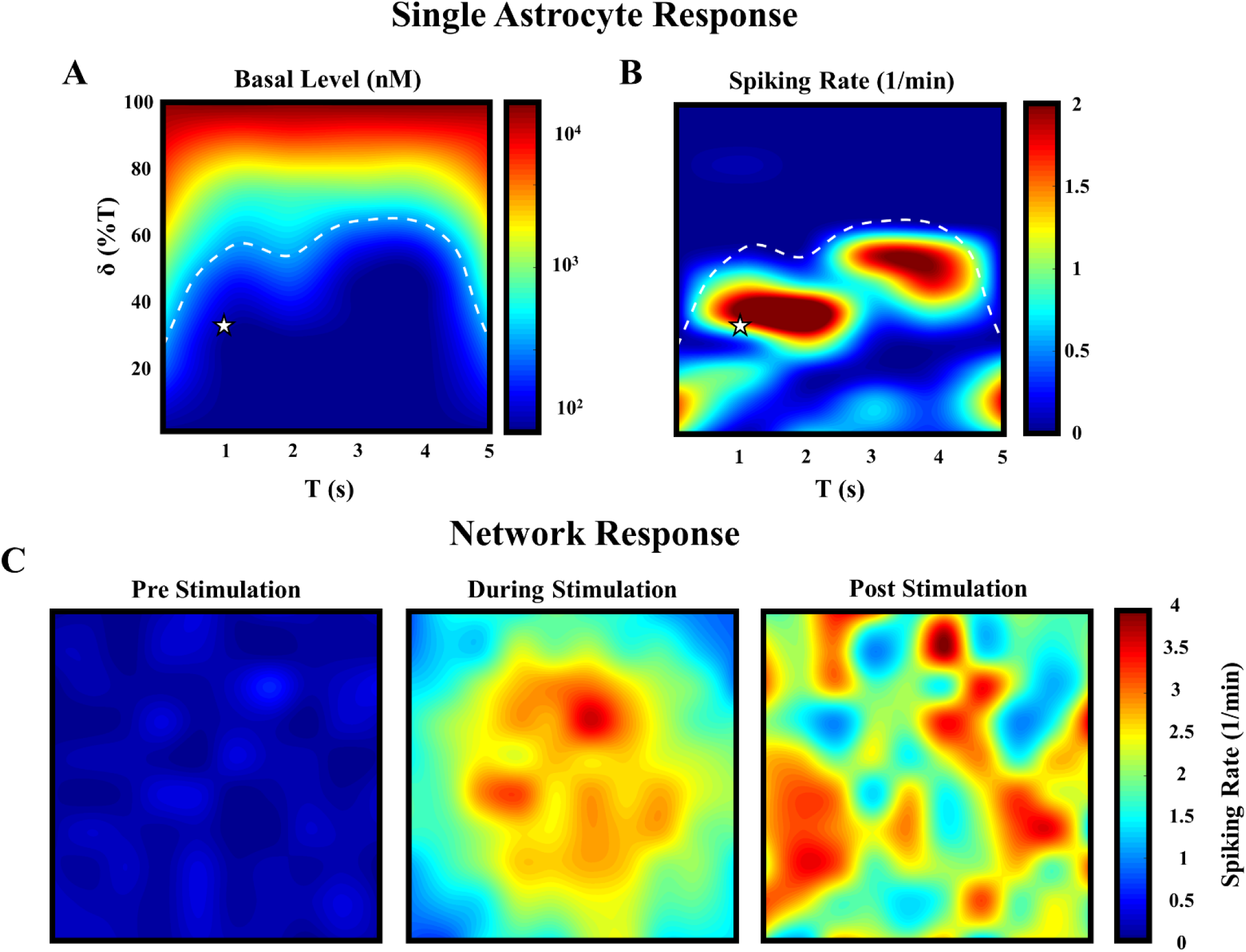
Response of ChRwt2 - expressing astrocytes to light stimulation. **A.** Heat map of the Ca^2+^ final basal level for various combinatorial windows of time duration (T) and pulse widths (δ; expressed as a percentage of T), expressed in the log scale. The physiological levels of Ca^2+^ basal level are indicated by the white dashed line. **B.** Heat map indicating spiking rate in the astrocyte for various combinatorial windows of T and δ, above the cutoff prominence chosen in Figure 3A. **C.** Heat maps indicating the mean spiking rate of astrocytes in the network - pre, during and post light stimulation (T = 1 s, δ = 0.3 s (30% of T), point denoted by star in **A** and **B**.

**Video S1. Movie of complete network-wide behavior of astrocytes to light stimulation.** In this video the stimulation window is marked in red. Parameters and stimulation specifics are as in Figure 5.

## Glossary

ChR: Channelrhodopsin;
PM: plasma membrane;
IP_3_: inositol trisphosphate;
IP_3_R: IP_3_ receptor;
CCE: capacitative calcium entry;
CICR: calcium induced calcium release;
SOC: store operated calcium channel;
ER: endoplasmic reticulum;
ATP: adenosine trisphosphate;
SERCA: sarcoplasmic reticulum Ca^2+^-ATPase;
PMCA: plasma membrane Ca^2+^ ATPase;
PLCδ: phospholipase C delta;
SCOs: spontaneous calcium oscillations;
LL: Local linearization;
LHS: Latin hypercube sampling;
PRCC: partial rank correlation coefficient

